# Mechanistic Basis for Microhomology Identification and Genome Scarring by Polymerase Theta

**DOI:** 10.1101/2019.12.20.882852

**Authors:** Juan Carvajal-Garcia, Jang-Eun Cho, Pablo Carvajal-Garcia, Wanjuan Feng, Richard D. Wood, Jeff Sekelsky, Gaorav P. Gupta, Steven A. Roberts, Dale A. Ramsden

**Author notes:** To whom correspondence should be addressed, Dale A. Ramsden.

## Abstract

DNA Polymerase Theta mediates an end joining pathway (TMEJ) that repairs chromosome breaks. It requires resection of broken ends to generate long, 3’ single stranded DNA tails, annealing of complementary sequence segments (microhomologies) in these tails, followed by microhomology-primed synthesis sufficient to resolve broken ends. The means by which microhomologies are identified is thus a critical step in this pathway, but is not understood. Here we show microhomologies are identified by a scanning mechanism initiated from the 3’ terminus and favoring bi-directional progression into flanking DNA, typically to a maximum of 15 nucleotides into each flank. Polymerase theta is frequently insufficiently processive to complete repair of breaks in microhomology-poor, AT-rich regions. Aborted synthesis leads to one or more additional rounds of microhomology search, annealing, and synthesis; this promotes complete repair in part because earlier rounds of synthesis generate microhomologies *de novo* that are sufficiently long that synthesis is more processive. Aborted rounds of synthesis are evident in characteristic genomic scars as insertions of 3-30 bp of sequence that is identical to flanking DNA (“templated” insertions). Templated insertions are present at higher levels in breast cancer genomes from patients with germline *BRCA1*/*2* mutations, consistent with an addiction to TMEJ in these cancers. Our work thus describes the mechanism for microhomology identification, and shows both how it mitigates limitations implicit in the microhomology requirement, and generates distinctive genomic scars associated with pathogenic genome instability.

## Introduction

DNA Double Strand Breaks (DSBs) in the chromosome are generated spontaneously, after exposure of cells to exogenous agents (e.g. ionizing radiation), and are induced during meiosis or development of the adaptive immune response (Chapman, Taylor and Boulton, 2012). DSBs are also generated by nucleases, especially Cas9, as intermediates in genome engineering. They are usually repaired by the Non-Homologous End Joining (NHEJ) pathway, which joins the two ends of a break with minimal processing (Waters *et al*., 2014), or the Homologous Recombination (HR) pathway, which uses another DNA molecule as a template for repair (Jasin and Rothstein, 2013). Impairment of either of these repair pathways – especially impairment in HR due to deficiency in *BRCA1* or *BRCA2* – leads to genome instability, which can cause cell death or cancer (Venkitaraman, 2002).

Another DSB repair pathway is defined by its requirement for DNA Polymerase Theta (Pol θ, gene name *POLQ*) (Chan, Yu and McVey, 2010; Roerink, Schendel and Tijsterman, 2014; Yousefzadeh *et al*., 2014), and has been termed Theta-Mediated End Joining (TMEJ). TMEJ overlaps with previously defined Alternative Non-Homologous End Joining (Alt-NHEJ) and Microhomology Mediated End Joining (MMEJ) pathways (Boulton and Jackson, 1996; Kabotyanski *et al*., 1998; Ma *et al*., 2003), though the extent these definitions overlap is not clear. In mammals, TMEJ is both more frequent and essential for viability in cells deficient in NHEJ (Wyatt *et al*., 2016; Saito, Maeda and Adachi, 2017; Zelensky *et al*., 2017) or HR (Ceccaldi *et al*., 2015; Mateos-Gomez *et al*., 2015; Feng *et al*., 2019). A specific requirement in *BRCA*-deficient contexts for Pol θ has identified this protein as a therapeutic target in *BRCA*-deficient breast cancers (Higgins and Boulton, 2018). However, TMEJ mechanism is not well understood, and it is important to determine its role and relevance in NHEJ and HR proficient cells.

TMEJ and HR pathways engage a common intermediate, the 3’ ssDNA tails generated after resection of chromosome breaks (Yousefzadeh *et al*., 2014; Wyatt *et al*., 2016). Pol θ aligns these tails and anneals small 2-6 bp patches of complementary sequence (microhomologies), which is followed by removal of at least one nonhomologous tail, then microhomology-primed synthesis sufficient to resolve remaining gaps (Fig. 1A) (Yousefzadeh *et al*., 2014; Kent *et al*., 2015). Differing location and frequency of microhomologies at different break sites is thus expected to have an impact on pathway outcome. At a minimum, the extent of deletion associated with repair by TMEJ will be determined by the locations of microhomologies relative to the break site, and especially the means by which these microhomologies are identified. Breaks in microhomology-poor regions of the genome could also lead to impaired TMEJ activity, and consequently cell death or cancer-causing genome rearrangements.

**Fig. 1.**
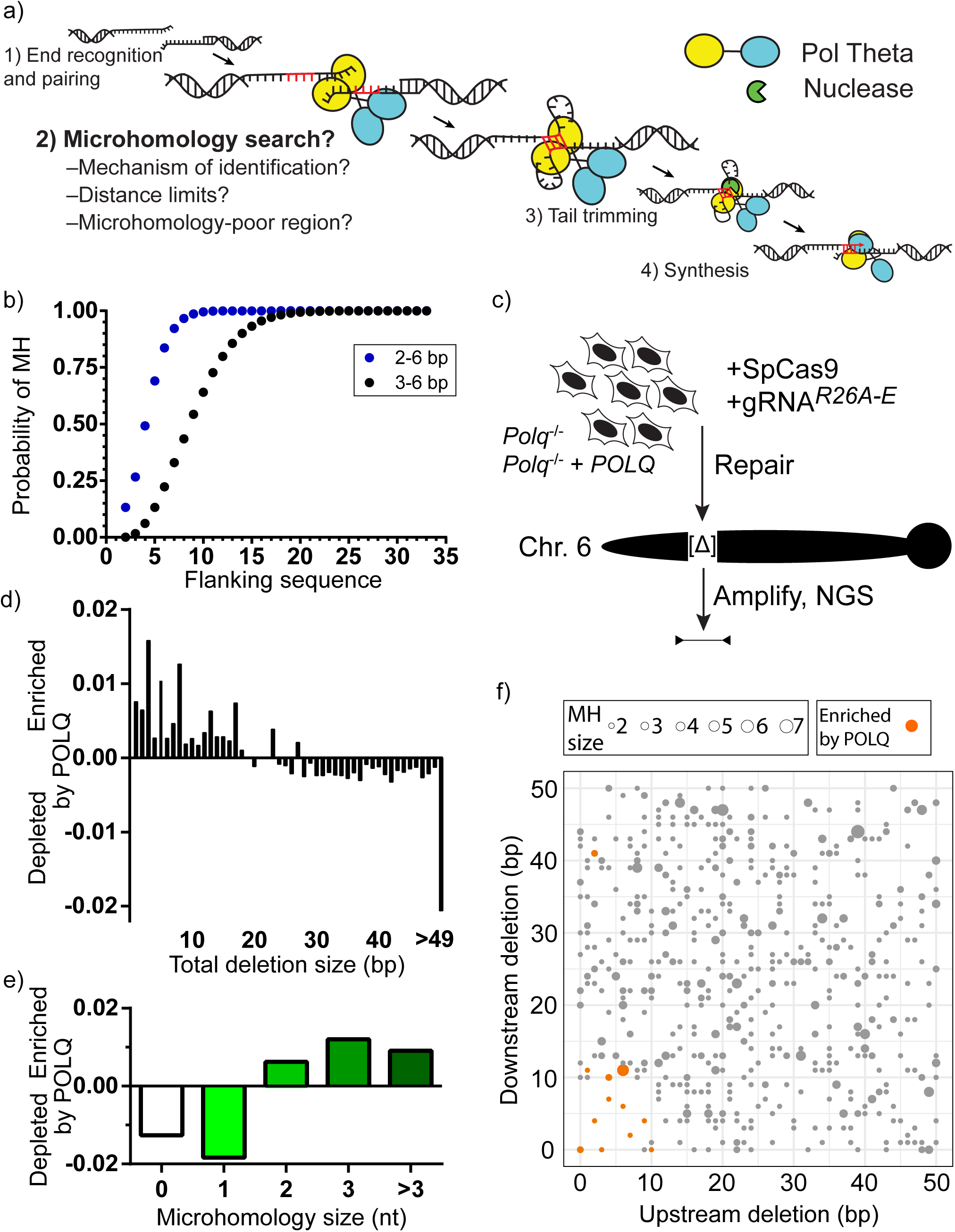
Characterization of Pol θ dependent deletions after a chromosome break. A) Steps required for TMEJ, emphasizing a critical role for microhomology identification. B) The probabilities of finding 2-6 bp (blue) or 3-6 bp (black) microhomologies (MH) were determined for sets of 100,000 randomly generated pairs of sequences of increasing size (flanking DNA sequence), from 2-33 nucleotides (nts). C) Cas9 targeted to 5 different break sites in the Rosa26 locus (R26A-E) were separately introduced into transformed mouse embryo fibroblasts (MEFs) from *Polq*^-/-^ deficient mice engineered to express human *POLQ* or not. Chromosome break repair products were recovered 24 hours later, amplified, and characterized by next generation sequencing (NGS). D) and E) The difference in the fraction of repair products with noted size of deletion (D) or microhomology (E) in *POLQ* expressing cells vs. *Polq^-/-^* was averaged across all 5 break sites tested. F) Filled circles denote the location of all microhomologies 2 bp or more relative to the break site for all 5 break sites, with microhomology size noted according to the size of the filled circle. Deletions enriched in cells expressing wt *POLQ* vs. *Polq*^-/-^ cells in triplicate experiments are shown in orange, and were identified using a two tailed t-test and the Benjamini-Hochberg procedure to adjust p-values for multiple comparisons, with a false discovery rate of 0.05.

**Fig. 2.**
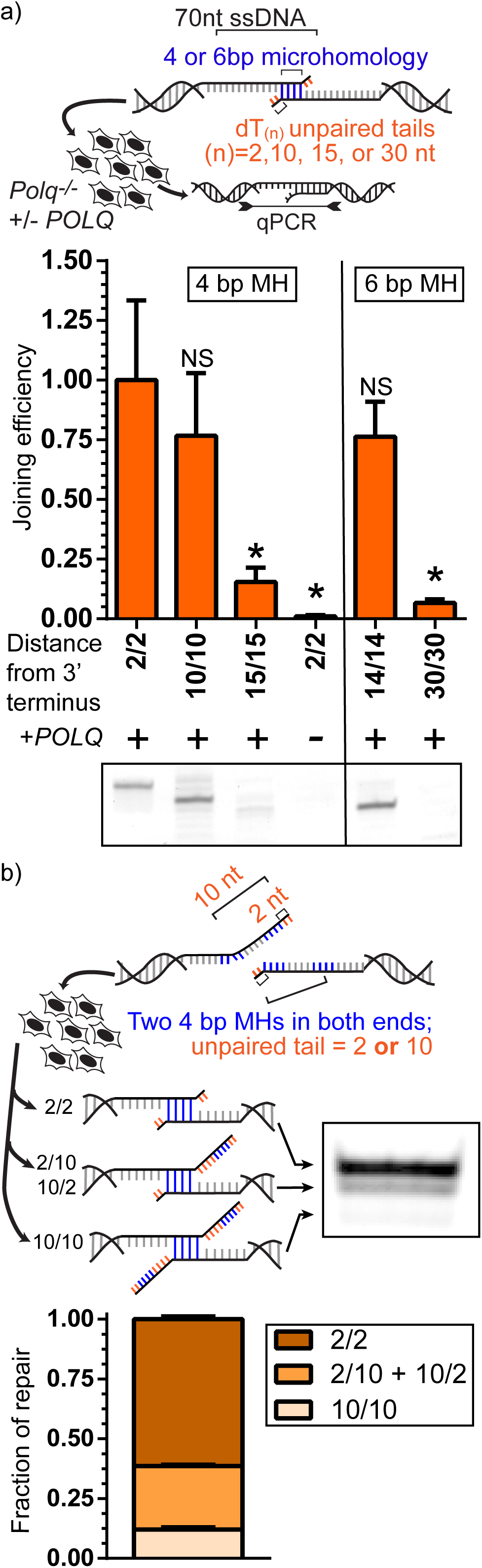
Mechanistic basis for microhomology usage by Pol θ. A) 691 bp dsDNA substrates with 70 nt 3’ ssDNA tails possessed microhomologies of 4 and 6 bp (4 bp MH, 6 bp MH) that were located in ssDNA tails 2, 10, 14, 15, and 30 nt from both head and tail 3’ termini (e.g. 2/2). Sequence 3’ of the microhomology was replaced with polyT(n) tracts. Substrates were introduced into the MEFs with and without *POLQ* expression described above. Head to tail end joining efficiencies in recovered DNA were determined by qPCR, and normalized to the joining efficiency observed using the 2/2 substrate. Statistical significance was assessed by one-way ANOVA with Bonferroni correction to account for multiple comparisons; ns, not significant; *, p <0.05. Experiments on substrates with 4 bp MH and 6 bp MH substrates were performed independently. Electrophoresis of a representative end point PCR is shown to confirm preferential usage of terminal 4 and 6 bp MHs for repair (bottom panel). B) Identical 4 bp MHs were located 2 nt and 10 nt from both head and tail 3’ termini and introduced into the cells described above. Products were amplified and characterized by electrophoresis (representative experiment shown); the mean relative amounts of noted species were determined for three independent experiments. Error bars denote the SEM.

Here we explore the basis for the microhomology identification step in TMEJ by systematically defining Pol θ-dependent repair products at a series of Cas9-induced chromosome breaks, where the break sites were chosen to possess varied density of break-site proximal microhomologies. We then explored the mechanistic requirements for TMEJ that are suggested by these results using an extrachromosomal substrate assay, as this latter strategy allows for unambiguous assessment of TMEJ activity on systematically varied substrates. Our results support a bi-directional scanning mechanism that mitigates deletion associated with TMEJ, as this scanning mechanism efficiently identifies microhomologies only when they are within 15 bp of either side of the break site. Break site locations within the genome that are depleted of microhomologies within 15 bp can nevertheless still be effectively repaired by TMEJ. In such contexts, TMEJ requires one or more cycles of aborted synthesis that generate long microhomologies *de novo*, which now better support processive synthesis. These latter products possess locally templated insertions that are highly characteristic of this pathway, and are consequently an effective biomarker for Pol θ/TMEJ activity in breast cancer genomes.

## Materials and Methods

### Cell lines

*Polq*^-/-^ Mouse Embryonic Fibroblasts (MEFs) were generated and immortalized with T antigen as described (Yousefzadeh *et al*., 2014) from *Polq*-null mice generated by conventional knock-out (Shima, Munroe and Schimenti, 2004) that were obtained from Jackson Laboratories and maintained on a C57BL/6J background. Pol θ was expressed by introducing wt *POLQ* human cDNA by lentiviral infection, with cells maintained in medium containing 4 μg/ml of puromycin. Cells were incubated at 37 °C, 5% CO2 and cultured in DMEM (Gibco) with 10% Fetal Bovine Serum (VWR Life Science Seradigm) and Penicillin (5 U/ml, Sigma). These lines and variants described below were confirmed to be free of mycoplasma contamination by a qPCR (Janetzko *et al*., 2014) with a detection limit below 10 genomes/1ml. Cell lines were additionally selected at random for third party validation of PCR results using Hoechst staining (Battaglia *et al*., 1994).

### Extrachromosomal assay

As has been described previously (Wyatt *et al*., 2016), extrachromosomal substrates consist of a 557 bp core DNA duplex ligated to head and tail caps with end structures that were varied as described in each figure. 75 ng of these substrates were electroporated into 200,000 cells with 1 ug of pMAX-GFP (Lonza) using the Neon system (Invitrogen) in a 10 ul tip with a 1,350 V, 30 ms pulse (three pooled electroporations formed a biological replicate) and incubated for 1 h. Cells were washed with Hank’s balanced saline solution, incubated with 25U of Benzonase (Sigma), DNA purified using the QIAamp DNA mini kit (QIAGEN), and products analyzed using SYBR green qPCR or a 30-cycle end point PCR. Each experiment consisted of three replicates of the above protocol. Quantification of gel bands was done with ImageQuant 8.1

### Chromosomal assay

Chromosomal DSB repair assays were performed using Cas9-gRNA RNP complexes assembled from Cas9 purified after overexpression in bacteria (Lin *et al*., 2014) (Addgene #69090), as well as annealed tracrRNA and target-specifying crRNA with stabilizing modifications (Alt-R, IDT). 7 pmol of Cas9 were incubated at room temperature with 8.4 pmol of annealed crRNA+tracrRNA for 30 mins, and electroporated into 200,000 cells with 32ng of pMAX-GFP as described above (three pooled electroporations formed a biological replicate). Cells were incubated for 24 hours and DNA was harvested with the QIAamp DNA mini kit (QIAGEN). Each experiment consisted of three biological replicates. DNA equivalent to 60,000 genomes was amplified for 24 cycles using primers purified by polyacrylamide gel electrophoresis (IDT) that included a 6 bp barcode, a spacer sequence of varying length (1-8 bp) to increase library diversity, and 21 (fwd primer) or 22 (rev primer) bp of Illumina adapter sequence. Amplicons were then purified using a 2% agarose (Lonza) gel and the QIAquick Gel Extraction Kit (QIAGEN), recovered DNA amplified with secondary NGS PCR primers for 5 cycles, and purified with AMPure XP beads (Beckman Coulter). Libraries were sequenced using a 300-cycle MiniSeq Mid Output Kit (R26A) or an iSeq 100 i1 Kit (R26B-E), including 20% of PhiX Control v3 DNA (Illumina). Data was analyzed using CLC Genomics Workbench 8 and Microsoft Excel.

### Statistical analysis

To identify microhomology associated deletions (MHD) significantly depleted in cells lacking *Polq*, we first identified the set of deletion products where each product was represented at a mean frequency greater than 1x10^-5^ in cells expressing *POLQ*. We then compared the frequencies for each product from 3 biological replicates for both *Polq*^-/-^ cells vs. *Polq*^-/-^ + *POLQ* cells using a two tailed t-test without sample pairing, and employed the Benjamini-Hochberg method with a false discovery rate of 0.05 to limit false positives that arise from making multiple comparisons. Calculations were performed in Microsoft Excel. Other experiments employed statistical tests as indicated in the figure legends using GraphPad Prism 8.

### High throughput sequencing junction characterization

Junctions were characterized by independently identifying the least-deleted 10 nucleotide match to sequence upstream of the break site, then the least-deleted 10 nucleotide match to sequence downstream of the break site. Microhomologies were defined as the overlap between upstream and downstream matches, and insertions were defined as non-matching sequences separating upstream and downstream matches. The location of microhomologies is defined as the distance relative to the break point after excluding microhomologous sequence.

For Figs. 3 and 4, insertions were characterized as templated direct repeats if the first 5 nucleotides of the inserted sequence could be mapped to sequence within 50 nt downstream of the break site (“downstream direct repeats”) or the last 5 nucleotides of the inserted sequence could be mapped to sequence 50 nt upstream of the break site (“upstream direct repeats”). Insertions were characterized as templated inverse repeats if the 5 last nucleotides of the inserted sequence could be mapped to the reverse complement of sequence within 50 nt downstream of the break (“right inverse repeats”) or if the 5 first nt of the inserted sequence could be mapped to the reverse complement of the 50 nt upstream of the break (“left inverse repeats”). We included cases of insertions < 5 nt as templated if additional templated insertions could be inferred due to involvement of 2° microhomologies in resolution (i.e. when sequence downstream of the insertion extended the identity that was detected in the proposed template to a total of 5 nt or more). We excluded insertions where the first inserted nucleotide was substituted, relative to the reference, but subsequent inserted sequence was identical to reference; such products could be identified in control experiments as substitutions made during sample amplification.

**Fig. 3.**
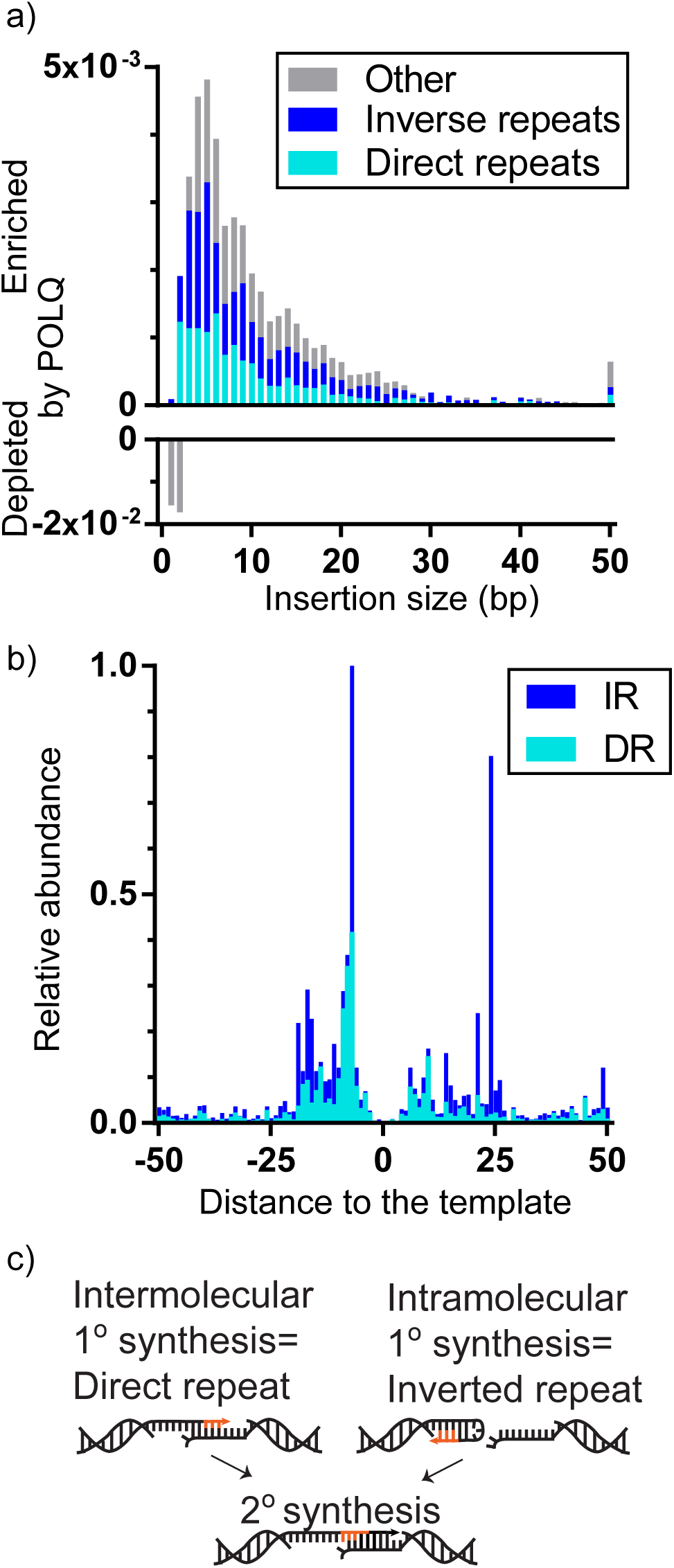
Pol θ generates locally templated insertions. A) The difference in the fraction of repair products with noted size of insertion in *POLQ* expressing cells vs. *Polq*^-/-^ was averaged across all 5 break sites. Insertions were defined as having 5 bp or more direct or inverse repeated sequence (DR, cyan; IR, blue) relative to flanking DNA (see also panel C). B) The relative abundance of insertions with 5 bp or more repeated sequence is plotted according to the location of the repeat within flanking DNA for both DR (cyan) and IR (blue) insertions. C) Model for generating direct repeat vs. inverted repeat templated insertions.

**Fig. 4.**
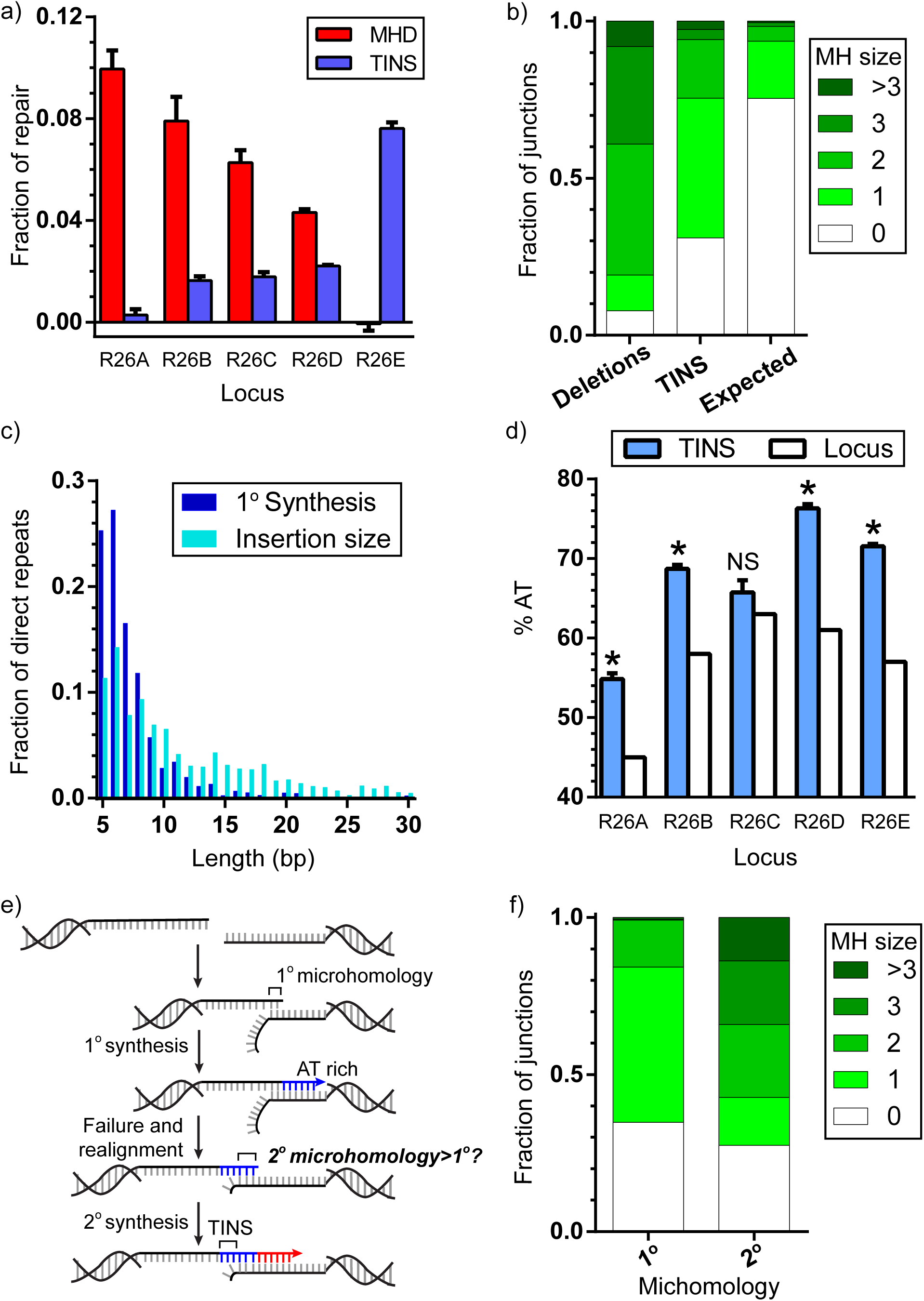
Characterization of TINS. A) Fraction of repair events enriched by *POLQ* expression that generated deletions with 2 bp or more of microhomology within 15 bp of the break site (MHD, red), or a templated insertion larger than 2 bp (TINS, blue), for each of the 5 break sites tested. Bar represents the mean and error bars SEM from 3 biological replicates. B) Average fraction of junctions associated with the indicated microhomology sizes in deletions significantly enriched by *POLQ* expression (Deletions), in insertions larger than 2 bp and directly repeated relative to flanking DNA (TINS), and as would be expected by chance, if microhomologies played no role (expected). C) Fraction of products with insertions of 5 bp or more sequence directly repeated relative to flanking DNA, comparing total length of synthesized nucleotides (insertion size; cyan) to the length of the first round of synthesis (1° synthesis; dark blue). D) Percent AT content in TINS defined as in C) (blue), compared to AT content in the 100 bp surrounding the break site for each locus (locus; white). Bar represents the mean and error bars SEM for 3 biological replicates. Statistical significance was assessed by a one-sample t test on each locus individually; ns, not significant; *, p<0.05. E) Model for the role of TINS in generating *de novo* (2°) microhomologies. F) Fraction of repair events associated with the indicated microhomology sizes for products with 4 or 5 bp of TINS in R26E, for 1° microhomologies vs. 2° microhomologies (see also panel E).

For Fig. 6 and S5, mutations from 569 whole genome sequenced breast cancers were obtained from the International Cancer Genome Consortium (ICGC) data portal (https://dcc.icgc.org/api/v1/download?fn=/release_26/Projects/BRCA-EU/simple_somatic_mutation.open.BRCA-EU.tsv.gz). Mutation lists were sorted by sample, chromosome and position prior to removing duplicate mutation entries. All insertion mutations and multiple nucleotide variants (MNV, which involve deletion of a sequence and insertion of a new, non-reference sequence) were extracted. For MNVs, the inserted sequences were compared to the deleted reference sequences to confirm accurate determination of the position the MNV event. For a small portion of MNVs, several nucleotides at either the 5’ or 3’ portions of the inserted sequences matched the sequence of the reference. These matching sequences were removed from the insertion sequence and the position of the MNV corrected prior to subsequent analysis. 100 nt of DNA sequence flanking each mutation was retrieved from the hg19 reference sequence for the human genome. Templated insertions (TINS) events among the breast cancer mutations were defined as above, except we considered only 30 nt of flanking DNA sequence as possible template. Breast cancer mutations were additionally stratified by *BRCA1* and/or *BRCA2* deficiency as determined by a germline mutation in either gene or hyper methylation as reported (Wen and Leong, 2019). Original data files describing the *BRCA1*/2 status of ICGC characterized tumors can be found at https://github.com/wenweixiong/BRCA2018. We obtained gene expression data for the evaluated tumors as log2 transformed fragments per kilobase per million reads (FPKM) RNA-seq from Supplementary Table 7 of (Nik-Zainal *et al*., 2016). We normalized *POLQ* log(2) FPKM counts according to housekeeping gene expression data as follows: for each of three housekeeping genes (*TBP*, *HPRT*, and *GAPDH*), we determined the difference between the log(2) FPKM counts for each tumor from the mean counts for the whole set, then averaged these three numbers for each tumor to determine normalization factors, which were then subtracted from the *POLQ* log(2) FPKM counts for each tumor.

**Fig. 5.**
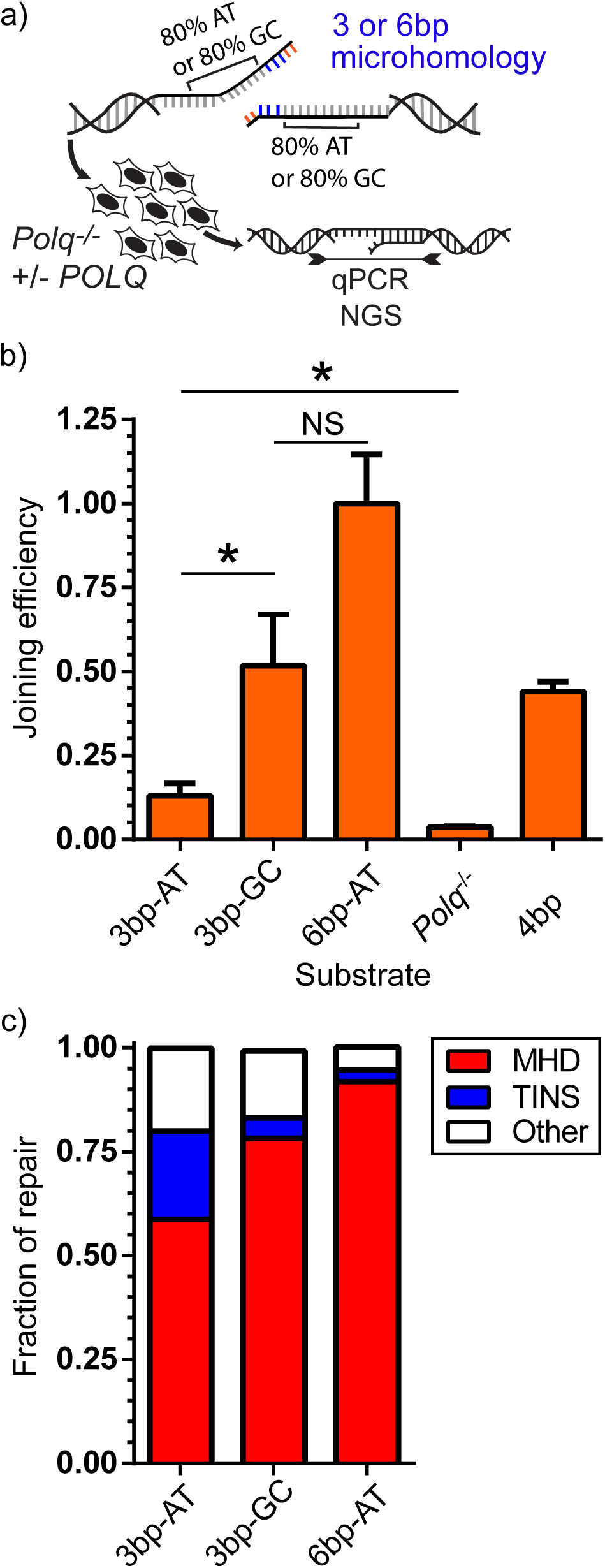
Frequency of TINS is dependent on both microhomology size and template AT content. A) TMEJ substrates possessed the same 3 bp microhomology followed by 20 bp of template that was either 80% AT (3 bp-AT) or 80% GC (3 bp-GC), or which possessed the 80% AT content template, but a longer 6 bp microhomology (6 bp-AT). The efficiency of cellular TMEJ (B) was determined for each substrate as in Figure 2A, and product structures (C) were characterized by next generation sequencing (NGS). B) Joining efficiency was determined by qPCR and normalized to results using the 6 bp-AT substrate, and compared to results using the original 2/2 4bp (4bp) substrate described in figure 2A. Bars represents the mean and error bars the standard error of the mean (SEM) from 3 biological replicates. Statistical significance was assessed by one-way ANOVA with Bonferroni correction to account for multiple comparisons; ns, not significant; *, p<0.05. C) TMEJ products were amplified and characterized by sequencing as MHD if possessing deletions at 2 bp or more of microhomology, or TINS if products contain 5 bp or more of templated (directly repeated) synthesis. Other represents products inconsistent with MHD or TINS.

**Fig. 6.**
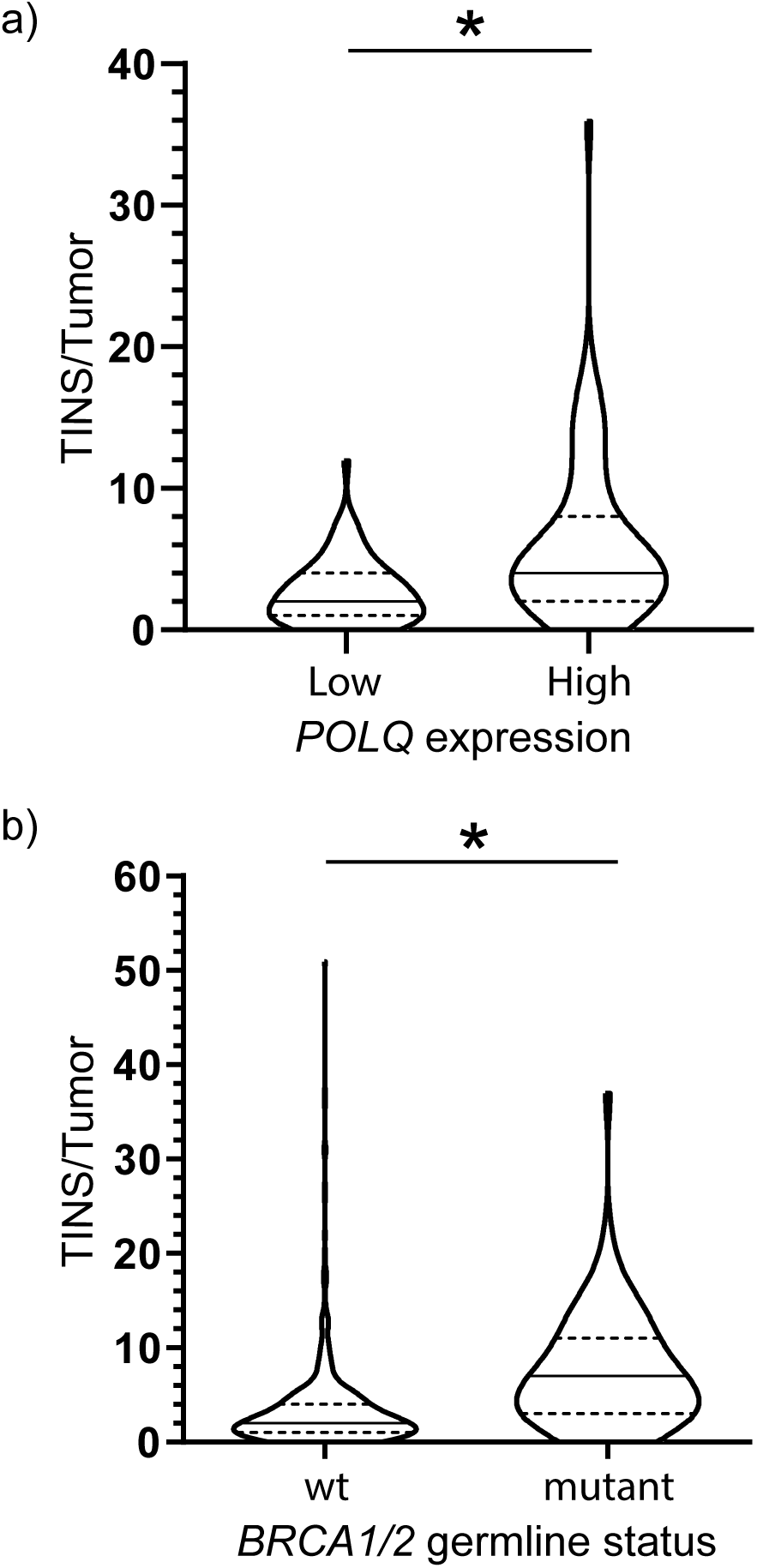
TINS are increased in tumors with high *POLQ* expression and with *BRCA* mutations. A) and B) Number of insertions with templated synthesis 5 bp or more per tumor (TINS/tumor, defined as in figure 4A) for breast cancer genomes previously sequenced by ICGC were determined according to differing *POLQ* mRNA expression level (A) or *BRCA1* or *BRCA2* germline status (B). A) *POLQ* expression in fragments per kilobase of transcript per million was normalized based on *HPRT1*, *TBT* and *GAPDH* expression for 261 breast cancers. Tumors were categorized as expressing low (*POLQ* Low; bottom quartile) or high levels of *POLQ* mRNA (*POLQ* High; top quartile). Statistical significance was assessed with a two-tailed Mann-Whitney test; *, p<0.05. B) Breast cancers were categorized as having germline *BRCA1* or *BRCA2* mutations (mutant, 75 cancers) or not (wt, 494 cancers).

## Results

### Characterization of Pol θ mediated deletions after a chromosomal break

Prior studies indicate efficient Pol θ mediated synthesis activity requires microhomologies of 3 bp or more (Kent *et al*., 2015; Wyatt *et al*., 2016; He and Yang, 2018). To assess how this requirement could impact repair we employed *in silico* modeling to determine the likelihood of finding such microhomologies as a function of increasing distance from the break site. The frequency of a 3 bp or more microhomology in a set of random pairs of break site flanking sequences is 64% when considering 10 bp of flanking sequence, and 93% when considering 15 bp. Inclusion of 2 bp microhomologies is sufficient to increase the frequency within 10 bp to 99% (Fig. 1B), though as discussed below, 2 bp microhomologies are less able to promote processive repair synthesis by Pol θ.

To address how these limitations impact TMEJ, we focused on 5 break sites (termed R26A through R26E) within a 7 kb region that varied according to the density and length of microhomologies near the break site (Fig. S1A). In particular, two closely located sites (within 500 bp) are unusually rich (R26A) vs. unusually poor (R26E) in terms of the availability of microhomologies within 15 bp.

We generated chromosome breaks at each of the 5 break sites by direct introduction of appropriately targeted *S. pyogenes* Cas9 ribo-nucleoprotein complexes into T antigen transformed mouse embryo fibroblasts (MEFs). We harvested cells 24 hours later, as this is sufficient time to accumulate repair products with insertions and deletions at the majority of chromosomes, while still mitigating the contribution of repair after excessive non-specific degradation of DNA ends. We amplified products without a phenotype-based screen, to ensure product spectra is limited only by whether the products can be amplified (i.e. retains primer sequences), then characterized products by next generation sequencing (Fig. 1C).

We first analyzed simple deletion products; i.e., deletions of flanking DNA without inserted sequences. We systematically identified those products enriched in human *POLQ* expressing MEFs (*Polq*^-/-^ MEFs complemented by expression of *POLQ* cDNA), relative to the *Polq*^-/-^ deficient isogenic parental cell line (Yousefzadeh *et al*., 2014). Highly enriched products consisted exclusively of deletions <30 bp (Fig. 1D) that were associated with 2-6 bp microhomologies (microhomology associated deletions; MHD) (Fig. 1E). The microhomologies chosen are also largely restricted to those within 10-15 bp of either side of the break site (Fig. 1F, Table S1). Moreover, nearly all microhomologies present less than 10 bp from termini (7/8) are significantly enriched (Fig. 1F). In accord with a restriction to 15 bp flanking the break site, we observed no significantly enriched MHD in repair products recovered from breaks at R26E, the site with few break-site proximal microhomologies (Fig. S1A, Table S1). More broadly, MHD associated with large deletions (>15 bp from either end) are actually suppressed by Pol θ/TMEJ for all 5 break sites tested, including R26E (Fig. S1B). TMEJ thus promotes more accurate repair.

### Mechanism of microhomology search

We addressed the mechanistic basis for microhomology choice described above by employing a cellular TMEJ assay that allows for systematic variation of the substrate. This assay employs extrachromosomal “pre-resected” substrates, consisting of double stranded DNA fragments with ends that have 70 nt single stranded DNA 3’ tails. The pre-resected tails block engagement of KU-dependent NHEJ, thus joining relies exclusively on Pol θ for efficient repair (joining efficiency reduced over 10-fold in *Polq* deficient cells; e.g. Fig. 2A) (Yousefzadeh *et al*., 2014; Wyatt *et al*., 2016).

We first assessed if a distance restriction on the ability of Pol θ to identify microhomologies helped explain the chromosomal repair results described above. We compared activity on substrates where a defined microhomology was present at increasing distance from the two 3’ termini, and replaced sequence downstream (3’) of the embedded microhomology with poly(dT) tracts, to ensure the absence of even trivial alternate microhomologies closer to the 3’ terminus. These substrates were introduced into the same isogenic MEFs with or without *POLQ* expression as described above, and repair assessed by quantifying the efficiency of repair, as well as by characterizing repair product structure (Fig. 2A).

TMEJ was similarly efficient when a 4 nt microhomology was embedded 2 or 10 nt from both 3’ termini, but was only 15% of maximal levels when the microhomology was embedded 15 nt (identified as 2/2, 10/10 and 15/15; distances refer to the size of nonhomologous tail after microhomology alignment) (Fig. 2A). TMEJ was also reduced if only one copy of the microhomology was located 15 nt distal to the break (2/15; Fig. S2A), in accord with rare use of asymmetrically located microhomologies during chromosomal TMEJ (only 1 of 5 within 10 bp; Fig. 1F). These observations exclude a mechanism where one end is fixed, and search proceeds inward from the other (i.e. the microhomology search typically progresses inward from both termini, rather than only one).

A larger 6 bp microhomology allowed for partial rescue of repair efficiency when located more than 10 bp away (14/14; 75%, relative to 2/2), but repair was abolished for even this larger microhomology when it was located 30 bp distal from both 3’ termini (Fig. 2A). Therefore, the limitation of Pol θ-dependent chromosomal MHD to within 15 bp of break sites described above (Fig. 1F) is at least in part due to a reduced ability of Pol θ to use microhomologies further away (Fig. 2A).

Microhomologies that are 2 nt vs 10 nt from both 3’ termini are used with similar efficiencies when they are the most 3’ proximal microhomologies present. We considered next whether they are also functionally equivalent when in competition. We designed a single TMEJ substrate where both ssDNA tails had two copies of the same microhomology, with one copy located 2 nt from the 3’ terminus, and the other copy 10 nt from the 3’ terminus (2/2+10/10) (Fig. 2B). Strikingly, the most proximal pair of microhomologies (2/2) was employed 5 times as frequently as any of the more distally located microhomologies (10/10, as well as 10/2 and 2/10) (Fig. 2B). Analysis of chromosomal data similarly reflected a strong preference for the more 3’ proximal of two equivalent microhomologies (Fig. S2B). These results are indicative of a mechanism that “scans” for available microhomologies, initiating from the 3’ terminus.

### Pol θ generates locally templated insertions

The experiments above focused on simple deletions (i.e. loss of flanking DNA without inserted sequences). We considered next the subset of chromosomal repair products that contain inserted sequence, whether such products contained deletion of flanking sequence or not. Insertion lengths varied between 1 and 157 bp. Pol θ increases the fraction of repair products with insertions over 2 bp, most clearly those between 3 and 6 bp (Fig. 3A). For most of these products, inserted sequences can be defined as repeated relative to nearby flanking DNA sequence (typically within 30 bp; Fig. 3B). These are best explained if the insertions are products of template-dependent synthesis (Templated insertions; TINS) (Yoshida *et al*., 1995; Jäger *et al*., 2000; Khodaverdian *et al*., 2017). Most of the remaining insertions enriched in *POLQ* expressing cells (grey in Fig. 3A) likely also employ a local template, but the length of the 1° synthesis tract was not sufficiently long to pass the 5-nucleotide minimum value we employed to exclude Pol θ-independent insertions. A small fraction (<0.1% of total repair) of insertions employed template distal to the break site, including other chromosomes. We identified two classes of TINS: synthesis of 5 bp or more of sequence that is directly repeated, relative to flanking DNA (direct repeats, DR), or synthesis of 5 bp or more of sequence that is the reverse complement of flanking DNA (Inverse Repeats, IR) (Fig. 3C). The former is consistent with a primary round of synthesis (1° synthesis) initiated from one broken, resected end, using the second resected end as a template (intermolecular synthesis), while the latter is consistent with 1° synthesis that was initiated from a hairpin (intramolecular synthesis).

The frequency of TINS varied widely across different break sites. Notably, TINS frequency was almost undetectable at R26A, the break site that is unusually rich in break proximal microhomologies (Fig. 4A). In contrast, TINS accounted for the only significant class of Pol θ-dependent events at R26E (Fig. 4A), where flanking sequence is poor in microhomologies.

Decreased availability of break site-proximal microhomologies thus correlates with both decreased MHD and increased TINS. Prior work argues that both MHD and TINS involve microhomology-mediated synthesis (Chan, Yu and McVey, 2010). However, MHD are generated after a single round of alignment-directed synthesis that was sufficiently processive to complete repair. By comparison, TINS are generated when the initial round of alignment-directed synthesis fails before repair is complete, and is followed by one or more additional rounds of alignment and synthesis (Fig. 3C). We sought to address next whether we can identify a mechanistic basis for failed synthesis.

We first confirmed that like MHD, the 1° round of synthesis in TINS favors alignment at microhomologies; the frequency of use of microhomologies 2 bp or more in this context is higher than would be expected by chance (25%, vs. 7%) (Fig. 4B). Preference for microhomologies is also sufficient to explain the non-random siting of synthesis initiation that is readily apparent in a map of the sources of templated insertions (Fig. 3B). However, microhomologies associated with TINS are generally shorter, relative to those associated with MHD, where 93% of MHD are associated with microhomologies 2 bp or more (Fig. 4B).

We further probed this correlation using specific microhomologies. We identified three different microhomologies that varied in size, but which were similarly located relative to the break site, then compared the relative frequencies of the two classes of events for each of the microhomologies (Fig. S3A). A long, 6 bp microhomology (AGTCTT) in R26A led to MHD 254-fold more often than TINS. An intermediate-sized 3 bp microhomology (TCC) in R26B lead to MHD 75-fold more frequently than TINS. Finally, a short 2 bp microhomology (AT) in R26E generated MHD only 1.66-fold more frequently than TINS. A given microhomology consequently contributes to both classes of Pol θ dependent events, but the extent that MHD is favored over TINS decreases as the size of the initial microhomology decreases. We conclude the probability of an aborted round of synthesis (and resolution with TINS) is partly reliant on instability in the alignment of the 1° microhomology. Two characteristics of TINS argue the processivity of synthesis after initial alignment also has a critical role in determining whether this round of synthesis aborts. 1° synthesis tracts in insertions are short – rarely more than 10 bp (Fig. 4C). Longer insertions, while present (insertion size, Fig. 4C), involve more than one round of templated insertions (e.g. Fig. S3B). In addition, the AT content of TINS was enriched relative to flanking DNA for all 5 break sites tested (Fig. 4D). This presumably reflects less effective stabilization of alignments involving synthesis through AT rich flanking sequence (relative to GC rich templates), more frequent failure, and a requirement for a second alignment and round of synthesis.

Prior work has noted how unsuccessful rounds of synthesis in TINS can generate microhomologies *de novo* (2° microhomologies), and these 2° microhomologies can be employed in alignments that prime the next round of synthesis (Fig. 4E) (Chan, Yu and McVey, 2010; Yu and McVey, 2010). We show here that these 2° microhomologies were typically longer (57% >1 bp) than the initial, failed 1° microhomology driven alignment (16% >1 bp) (Fig. 4F). TINS is thus effectively an adaptive mechanism – in microhomology-poor regions it generated new, more stable microhomologies that increased the likelihood of successful repair.

We employed extrachromosomal substrates to directly test if TINS are a function of both reduced size of 1° microhomology, as well as the AT content of the template flanking the microhomology. We assessed the importance of flanking DNA by generating two substrates with the same short, 3 bp microhomology (TAG), but highly divergent levels of flanking AT content; a substrate with 80% AT content downstream of the microhomology, vs. a substrate with 80% GC content downstream of the microhomology (Fig. 5A). When flanking DNA was AT rich, joining was 25% as efficient as when flanking DNA was GC rich (Fig. 5B), and repair products generated using the substrate with AT rich flanking DNA more often possessed TINS (Fig. 5C). These results are consistent with a critical role for ongoing synthesis in stabilizing a given alignment, and enabling productive repair. We further used a variant of the substrate with AT rich flanking DNA with a longer microhomology (6 bp) (Fig. 5A). The longer 1° microhomology was sufficient to promote higher joining efficiencies (Fig. 5B), as well as lower frequencies of TINS (Fig. 5C), relative to both of the substrates (AT rich and GC rich) with 3 bp microhomologies.

### TINS genomic scars are a biomarker for increased TMEJ activity in *BRCA* mutated cancers

MHD are often – though not always (see e.g. Fig. 4A) – the most frequent Pol θ-dependent repair products for a given chromosome break, and serve as a biomarker for TMEJ activity (Feng *et al*., 2019). However, some MHD are favored during NHEJ (Fig. S4A), while yet others can be suppressed by Pol θ (Fig. S1B). By comparison, TINS are a more characteristic product of Pol θ-dependent repair (Fig. S4B), and consequently has been proposed as a more effective biomarker for Pol θ/TMEJ activity (Schimmel *et al*., 2019). To address this possibility, we assessed whether it was possible to correlate the frequency of TINS with *POLQ* expression levels in the genomes of breast cancers previously sequenced by ICGC (Nik-Zainal *et al*., 2016) and made publicly available at https://dcc.icgc.org/releases/release_26/Projects/BRCA-EU. We determined that TINS were higher in the 65 cancers with high *POLQ* expression (top quartile), relative to the 66 cancers with low *POLQ* expression (bottom quartile) (p<0.0001, Mann-Whitney test) (Fig. 6a). Notably, we observe a similar correlation of TINS with *POLQ* expression (p=0.0002, Mann-Whitney) after excluding cancers with germline mutations in *BRCA1* or *BRCA2*, a possible confounding issue (Fig. S4C) (discussed next).

Pol θ is required for viability in cancer cell lines deficient in *BRCA1*/*2* (Ceccaldi *et al*., 2015; Mateos-Gomez *et al*., 2015), and higher levels of *POLQ* expression are observed in *BRCA1*/*2* mutated breast cancers (Fig. S4D) (Higgins *et al*., 2010; Lemée *et al*., 2010; Ceccaldi *et al*., 2015). We show here that the genomes from the 75 breast cancer patients with germline *BRCA* mutations had a median of 7 TINS/genome, a much higher frequency than that observed in the remaining patients with no germline *BRCA* mutations (2 TINS/genome) (p<0.0001, Mann-Whitney test) (Fig. 6B). The ability to correlate *BRCA1*/*2* deficiency with increased frequency of TINS, a biomarker more directly reflective of Pol θ/TMEJ activity than expression, provides further support that addiction to TMEJ is broadly associated with *BRCA1*/*2* deficient cancers.

## Discussion

### Mechanistic basis for microhomology identification

TMEJ identifies microhomologies through a scanning mechanism that is initiated from the 3’ terminus, and favors break-proximal, 2 bp or larger microhomologies (Fig. 2B). Increased size of microhomology can modestly extend the distance searched beyond the 15 bp limit described above (Fig. 2A), and also impacts the extent proximal microhomologies are favored. Asymmetrically located pairs of microhomologies are used less efficiently than a symmetrically located pair of microhomologies (e.g. 10/10 vs. 2/15 substrates, 2A, S2A, or Fig. 1F), implying scanning is typically bi-directional.

Past work from our group and others has determined that in some contexts, Pol θ-dependent repair can include microhomologies more distal than 15 bp from 3’ termini. These contexts include cells deficient in regulation of end resection (cells defective in *KU* or *53BP1*) or wild type cells after a much longer recovery period than is used here (24 hours) (Wyatt *et al*., 2016; Schimmel *et al*., 2017; Feng *et al*., 2019). Given our results with pre-resected substrates, we suggest this second class of Pol θ-dependent products reflects loss of 3’ terminal ssDNA before Pol θ could be engaged, rather than a fundamental change in the preference of Pol θ for break proximal microhomologies. Notably, there is little mechanistic rationale for use of microhomologies >15 bp from either side of the break site, since i) for break sites rich in microhomologies in this region, the identification of these microhomologies is both efficient (Fig. 1F) and strongly favors the most break-proximal microhomologies (Fig. 2B), and ii) for the 7% of break sites without a break-proximal microhomology 3 bp or more (Fig. 1B), iterative rounds of synthesis can generate microhomologies *de novo* that are now longer, and which support processive synthesis and successful repair (Fig. 4).

### Mechanism of template-dependent insertions

Repair by simple Pol θ-dependent MHD is unlikely to be efficient at the 7% of break sites that do not have microhomologies 3 bp or more within 15 nt of the break site, and instead generates near-compensatory levels of Pol θ-dependent products with locally templated insertions (TINS; Fig. 4A). We show that both classes of Pol θ mediated repair – MHD and TINS – are products of synthesis initiated from primers annealed to template at sites of short complementary sequence. TINS reflect instances of alignment and synthesis where synthesis was not sufficiently processive to allow for complete repair. Such failures are primarily due to alignments that rely on short, <3 bp microhomologies or no microhomology, consistent with *in vitro* data emphasizing the importance of alignments of larger microhomologies for processive synthesis (He and Yang, 2018; Black *et al*., 2019). However, even after synthesis is initiated, there is frequent failure if synthesis proceeds through AT-rich regions (Fig. 4D, Fig. 5). Importantly, the initial aborted round of synthesis often generates a new, larger microhomology (Fig. 4F). This explains how TINS is compensatory in microhomology-poor, AT-rich regions – it is effectively an adaptive mechanism, as Pol θ can generate new microhomologies that are sufficiently stable to sustain processive synthesis and successful repair. We show that TINS is a specific marker for TMEJ activity (Fig. S4B). They are present at a much higher frequency in a panel of 75 breast cancers with germline mutations in *BRCA1*/*2* (Fig. 6B), consistent with a requirement for TMEJ for viability of several *BRCA1*/*2* defective cancer cell lines (Ceccaldi *et al*., 2015; Mateos-Gomez *et al*., 2015). Indeed, Pol θ is required for viability in the context of a wide variety of DNA damage response defects, and thus is as an attractive target for therapy in as many as 30% of all breast tumors (Higgins and Boulton, 2018; Feng *et al*., 2019). Assessment of TINS in tumor genomes thus holds promise as a biomarker for deciding when TMEJ should be targeted for cancer therapy.

### Relationship of TMEJ to other end-joining repair

We have defined the outcomes of TMEJ as i) those deletion products associated with microhomologies 2 bp or more that are located within 15 bp of either side of the break site, as well as ii) products with 5 bp or more of synthesis that results in an insertion, and the synthesis employs template within 50 bp of either side of the break site.

Loss of Pol θ does not typically lead to compensatory increases in the accurate, <5 bp deletion products that can be clearly linked to NHEJ (Fig. 7A) (Schimmel *et al*., 2017). Instead, repair is re-channeled to “other” products with larger deletions (>15 bp from either side of the break site; Fig. 7A) that also favor microhomologies, though to a lesser degree than TMEJ (Fig. 7B). The contribution of TMEJ to overall repair also varies little over the 5 break sites tested – an average of 8.4+/-1.5% (standard deviation) – despite wide variation in the availability of microhomologies. Taken together, our results imply 5-10% of Cas9-induced DSBs preferentially engage TMEJ and not NHEJ, likely because these DSB ends have been resected (Fig. 7C). Less accurate end joining products that are enriched in Pol θ deficient cells may at least in part reflect clipping of 3’ ssDNA tails generated by resection (e.g. by Artemis) (Wyatt *et al*., 2016; Shibata *et al*., 2017), followed by NHEJ or yet-undefined alternate end joining mechanisms.

**Fig. 7.**
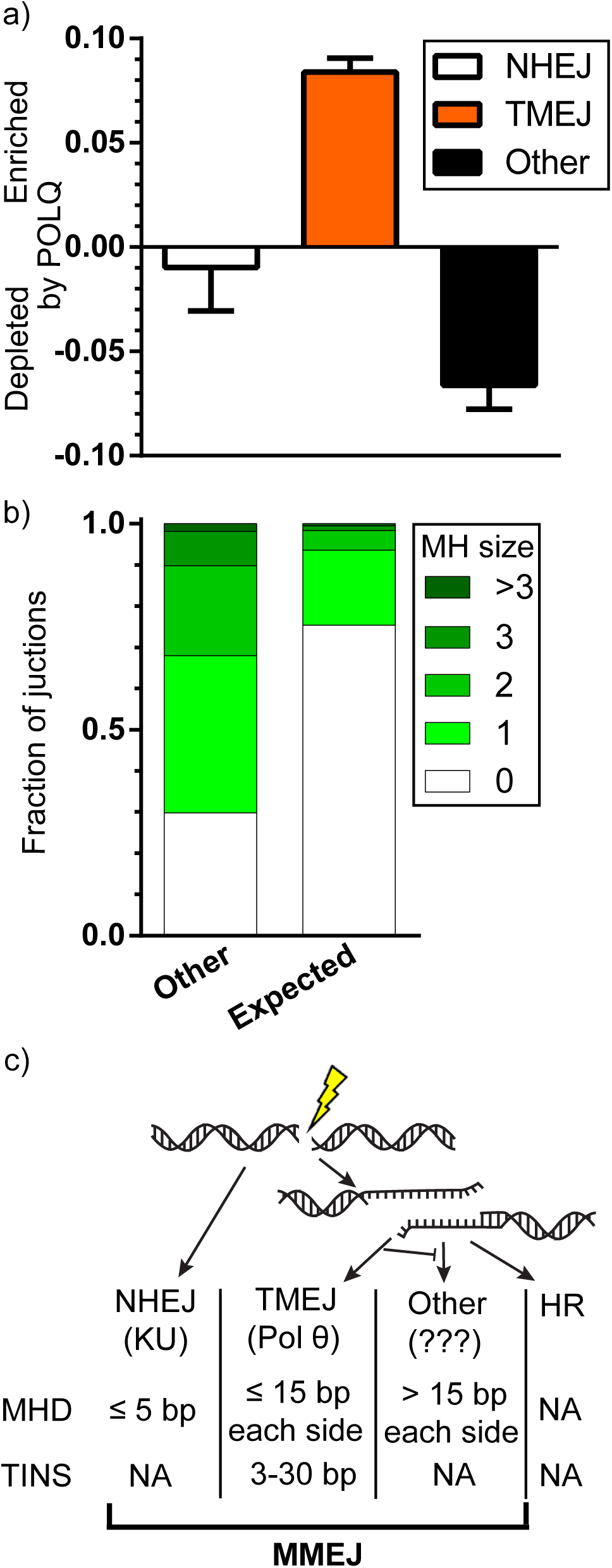
TMEJ promotes genomic stability. A) The extent of enrichment of repair product classes upon *POLQ* expression, averaged across all 5 break sites. Products were classified as NHEJ (white bar) if deleted sequence was less than 5 bp and possessed no or 1 bp of microhomology, as TMEJ (orange bar) if deleted sequence was less than 15 from either end and possessed 2 bp or more of microhomology, or had templated insertions larger than 2 bp, and as “other” (black bar) if deleted sequence from either flank exceed 15 bp. Bar represents the mean and error bars SEM from 3 biological replicates. B) Average fraction of repair events associated with the indicated microhomology sizes in deletions larger than 15 bp from both sides of the break (other) or as would be expected by chance, if microhomologies played no role (expected). C) Following a double strand break (DSB), ends are either ligated through the NHEJ pathway (dependent on the KU heterodimer) or resected to form 3’ ssDNA tails. These are substrates for TMEJ (dependent on Pol θ), HR, or a third, undescribed form of end joining that also favors microhomologies. Different end-joining pathways have different mutational outcomes (MHD of the indicated sizes or TINS).

A role for microhomologies in mammalian end joining has long been clear (Roth and Wilson, 1986). Our work shows they have a variety of sources. Some microhomology associated deletions (MHD) are generated by both Pol θ/TMEJ and NHEJ (those with less than 5 bp of deletion; Fig. S4A), while others are primarily attributable to Pol θ/TMEJ (between 5-15 distal to either flank; Fig. 1F). Yet a third class (those suppressed by TMEJ in otherwise wild type cells, and more than 15 bp from either strand-break terminus; Other, Fig. 7A) has distinct and probably complex genetic requirements. MHD are thus generated by at least three genetically distinguishable mechanisms, highlighting a critical flaw in the frequent attribution of such products to a single microhomology mediated end joining pathway (“MMEJ”) (Fig. 7C).

Implicit to a requirement for microhomologies during end-joining is i) an associated loss of genomic information, and ii) potentially impaired repair in contexts where microhomologies are hard to find. We describe here how an elegant search mechanism largely overcomes these limitations in a manner that in most contexts – e.g. in cells proficient in canonical pathways, and which appropriately regulate end resection – least threatens genome stability. At the same time, cancer-causing deficiencies in BRCA genes leads to excessive engagement of TMEJ and “addiction” of cells to this pathway, which can help drive cancer progression.

## Author Contributions

Author contributions: J.C.G. and D.A.R. designed research; J.C.G., J.E.C. and W.F. performed research; P.C.G., R.D.W and S.A.R. contributed new reagents/analytic tools; J.C.G. and D.A.R. analyzed data; J.C.G., and D.A.R. wrote the paper; D.A.R., G.G. and J.S. supervised the research; D.A.R., G.G. and J.E.C. acquired funding.

## Acknowledgments

Our studies were supported by NCI grant CA222092 (D.A.R. and G.G.), DOD grant W81XWH-18-1-004 (D.A.R. and G.G.) and T32CA009156 (J.E.C.)

---

**Fig. S1.**
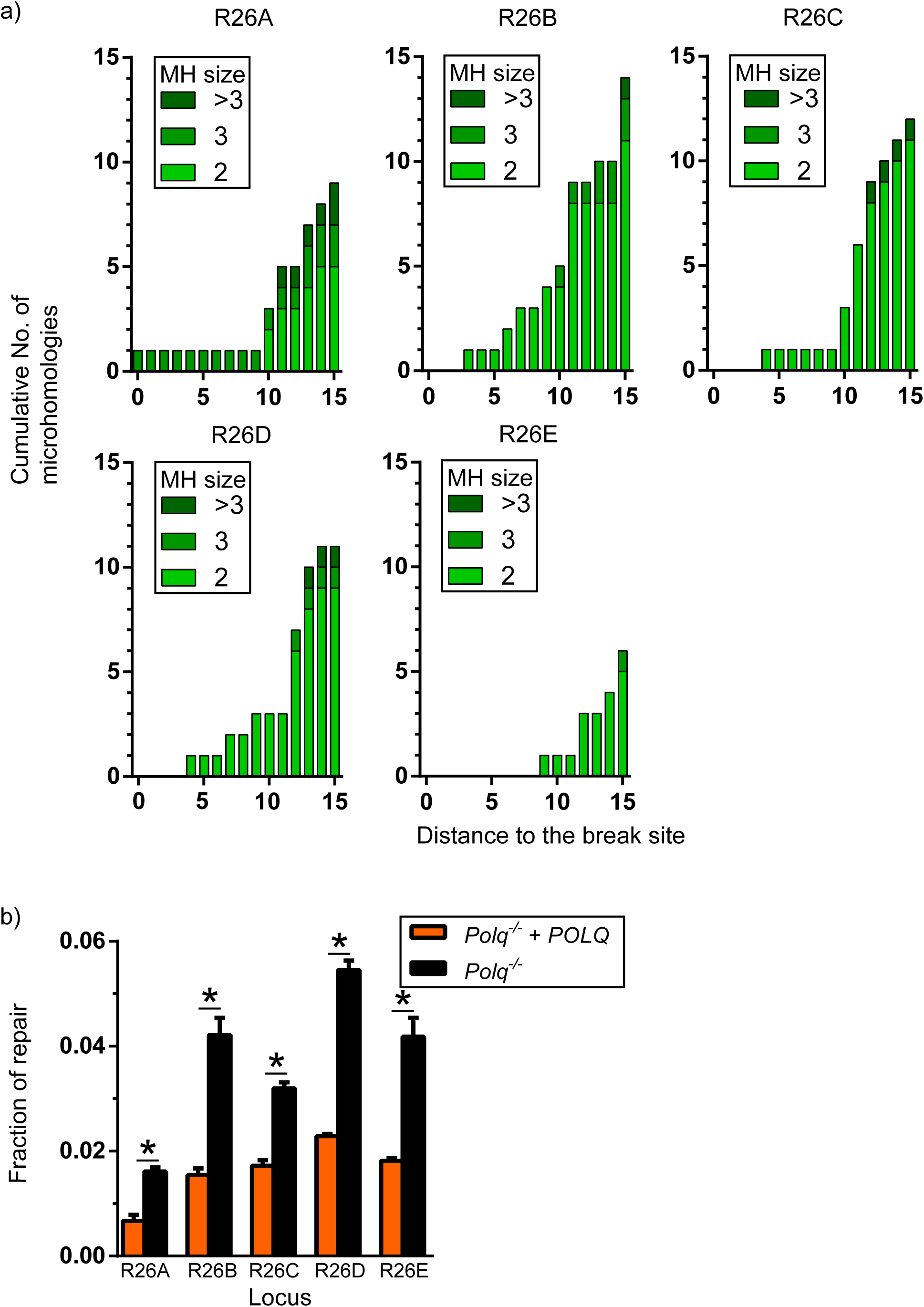
A) Cumulative number of microhomologies larger than 1 bp at the noted distance to the break site. Distance to the break site corresponds to the largest of the two distances (upstream and downstream) from the beginning of the microhomology to the break terminus. B) Fraction of repair corresponding to deletions with microhomologies 2 bp or more and located more than 15 nt from both sides of the break site in *Polq*^-/-^ cells that expressed POLQ (orange) or not (black). Bar represents the means and error bars SEM for 3 biological replicates. Statistical significance was assessed by one-way ANOVA with Bonferroni correction to account for multiple comparisons; *, p<0.05.

**Fig. S2.**
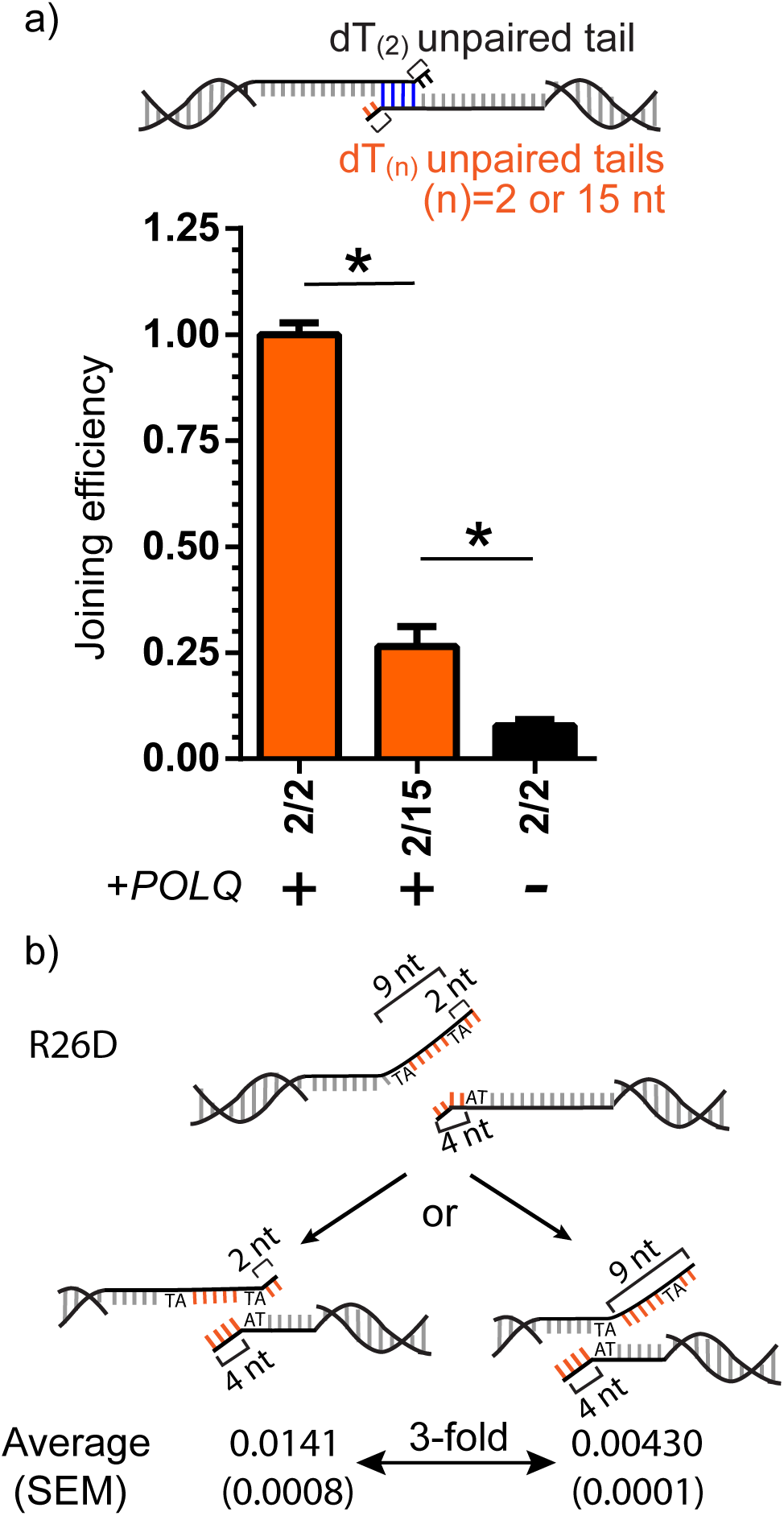
A) Joining efficiency was calculated as in Fig. 2 for a substrate with a microhomology located 2 nt away from the 3’ terminus for both head and tails ends (2/2), as well as a substrate with a microhomology 2 nt from the head 3’ terminus and 15 nt from the tail 3’ terminus (2/15). Bars represent the mean and error bars SEM for 3 biological replicates. Statistical significance was assessed by one-way ANOVA with Bonferroni correction to account for multiple comparisons B) Schematic of two Pol θ dependent MHD identified in chromosomal repair products at the break site R26D. The fraction of repair products enriched by *POLQ* expression was calculated as in Fig 1. SEM is shown in parenthesis for 3 biological replicates.

**Fig. S3.**
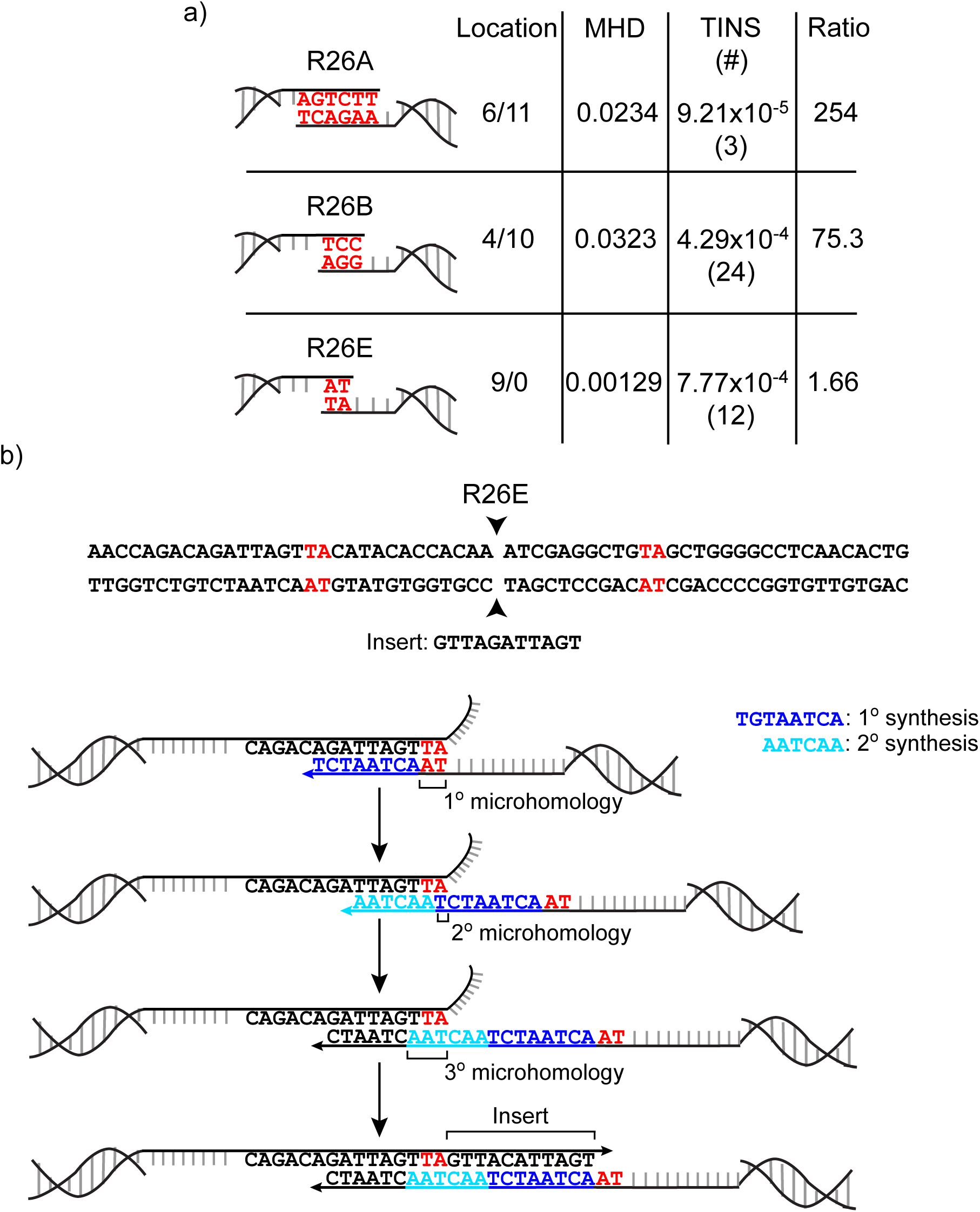
A) The fraction of repair products enriched by *POLQ* expression comparing MHD vs. TINS and number of different TINS identified (#) for each of three 1° microhomologies of differing length; a 6 bp AGTCTT microhomology in R26A, a 3 bp TCC microhomology in R26B, and a 2 bp AT microhomology in R26E). The location of the 1° microhomology with respect to the break site is indicated as upstream deletion/downstream deletions). B) Generation of a repair product consistent with three microhomology primed synthesis events. Primary (1°) microhomology is shown in red, 1° round of synthesis in dark blue, and secondary (2°) round of synthesis in cyan.

**Fig. S4.**
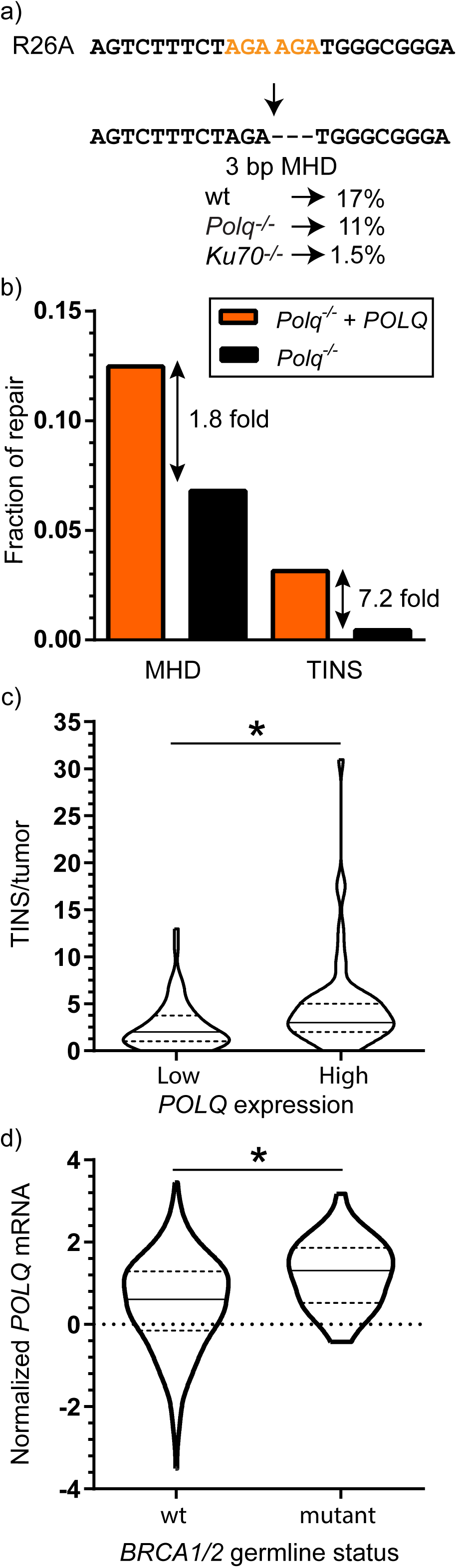
A) Structure of a repair event in R26A characterized as deletion of a terminal 3 bp microhomology (in orange), with the fraction of repair represented by this product noted for wild type, *Polq*^-/-^ and *Ku70*^-/-^ cells. Data for *Ku70*^-/-^ was obtained from (3). B) Fraction of repair corresponding to MHD and TINS averaged across the 5 break sites tested in cells expressing *POLQ* (orange) or not (black). The fold difference is shown and was calculated as the fraction in *POLQ* expressing cells divided by the fraction in parental *Polq*^-/-^ cells. C) The frequency of TINS/tumor genome was determined for tumors with high or low levels of *POLQ* expression determined as in Fig. 6a, except tumors with germline *BRCA* mutations were excluded. D) Levels of *POLQ* mRNA were normalized as in Fig. 6A, and compared in tumors with wild type (wt) germline *BRCA* genes vs. tumors with germline mutations in *BRCA* genes. Statistical significance was assessed with a two-tailed Mann-Whitney test, *p<0.05.

**Table S1:** Pol θ dependent repair events. Upstream and downstream deletion, microhomology size, and fraction of repair enriched by POLQ expression for all the repair events significantly enriched in cells expressing wt POLQ vs. Polq-/- in triplicate experiments. Statistical significance was identified using a two tailed t-test and the Benjamini-Hochberg procedure to adjust p values for multiple comparisons, with a false discovery rate of 0.05.

| Locus | Upstream deletion | Downstream deletion | Microhomology size | Fraction enriched by POLQ |
| --- | --- | --- | --- | --- |
| R26A | 0 | 0 | 3 | 0.057557 |
|  | 6 | 11 | 6 | 0.023439 |
|  | 10 | 0 | 2 | 0.006968 |
|  | 3 | 5 | 0 | 0.000259 |
| R26B | 4 | 10 | 3 | 0.032344 |
|  | 3 | 0 | 2 | 0.017555 |
|  | 1 | 11 | 2 | 0.008442 |
|  | 6 | 6 | 2 | 0.006746 |
|  | 3 | 4 | 1 | 0.004777 |
|  | 7 | 2 | 2 | 0.004391 |
|  | 8 | 4 | 1 | 0.00411 |
|  | 6 | 9 | 1 | 0.003844 |
|  | 2 | 6 | 0 | 0.001395 |
| R26C | 2 | 4 | 2 | 0.036814 |
|  | 9 | 9 | 0 | 0.000588 |
|  | 11 | 6 | 0 | 0.00058 |
|  | 3 | 7 | 0 | 0.000381 |
|  | 15 | 3 | 0 | 0.000305 |
| R26D | 4 | 7 | 2 | 0.021654 |
|  | 2 | 4 | 2 | 0.014109 |
|  | 1 | 4 | 1 | 0.008736 |
|  | 2 | 1 | 0 | 0.006699 |
|  | 0 | 1 | 0 | 0.005915 |
|  | 2 | 1 | 1 | 0.004951 |
|  | 9 | 4 | 2 | 0.004299 |
|  | 4 | 0 | 1 | 0.004095 |
|  | 8 | 4 | 1 | 0.002299 |
|  | 1 | 1 | 0 | 0.0018 |
|  | 9 | 1 | 0 | 0.001155 |
|  | 2 | 41 | 3 | 9.37E-05 |
| R26E | 6 | 6 | 0 | 0.00366 |

